# Combined effects of bird extinctions and introductions in oceanic islands: decreased functional diversity despite increased species richness

**DOI:** 10.1101/2021.10.11.463897

**Authors:** Filipa C. Soares, Ricardo F. de Lima, Jorge M. Palmeirim, Pedro Cardoso, Ana S. L. Rodrigues

**Author notes:** Corresponding author: Postal address: cE3c - Centre for Ecology, Evolution and Environmental Changes, Faculdade de Ciências, Universidade de Lisboa, 1749-016 Lisboa, Portugal.

## Abstract

**Aim:** We analyse the functional consequences of the changes in species composition resulting from extinctions and introductions on oceanic island bird assemblages. Specifically, we ask if introduced species have compensated the functional loss resulting from species extinctions.

**Location:** Seventy-four oceanic islands (>100 km^2^) in the Atlantic, Pacific and Indian Oceans.

**Time period:** Late Holocene.

**Major taxa studied:** Terrestrial and freshwater bird species.

**Methods:** We compiled a species list per island (extinct and extant, native and introduced), and then compiled traits per species. We used single-trait analyses to assess the effects of past species extinctions and introductions on functional composition. Then, we used probabilistic hypervolumes in trait space to calculate functional richness and evenness of original versus present avifaunas of each island (and net change), and to estimate functional originality of extinct and introduced species.

**Results:** The net effects of extinctions and introductions were: an increase in average species richness per island (alpha diversity), yet a decline in diversity across all islands (gamma diversity); an average increase in the prevalence of most functional traits (23 out of 35) yet an average decline functional richness and evenness, associated with the fact that extinct species were functionally more original (when compared to extant natives) than introduced species.

**Main conclusions:** Introduced species are on average offsetting (and even surpassing) the losses of extinct species per island in terms of species richness, and they are increasing the prevalence of most functional traits. However, they are not compensating the loss of functional richness due to extinctions. Current island bird assemblages are becoming functionally poorer, having lost original species and being composed of functionally more homogeneous species. This is likely to have cascading repercussions on the functioning of island ecosystems.

## 1. Introduction

Human activities are profoundly changing the distribution of species worldwide at an alarming pace: the composition of communities is being altered both through the local or global disappearance of some species and the introduction and expansion of others (McKinney & Lockwood, 1999). Oceanic islands are among the most threatened ecosystems and their assemblages have been largely shaped by the history of human occupation: compared to continents, islands tend to have higher extinction rates by being more sensitive to habitat modification and biological invasion (Loehle & Eschenbach, 2011; Whittaker et al., 2017; Russell & Kueffer, 2019). In particular, birds have suffered a high proportion of extinctions on islands (Sax & Gaines, 2008), which affected mostly large, flightless and ground-nesting species with specialized diets (e.g. nectivores and insectivores; Boyer & Jetz, 2014). Consequently, on many islands, these non-random extinctions led to a disproportionate loss of functional diversity (Boyer, 2008; Boyer & Jetz, 2014; Sobral et al., 2016), potentially causing a sharp decline in the variety of ecological functions provided by birds (e.g. Heinen et al., 2018) and ultimately affecting ecosystem functioning (Şekercioğlu et al., 2004; Sax & Gaines, 2008; Luck et al., 2012).

It is yet unclear if the introduction of species can functionally replace the loss of native species, although having been recently considered a fundamental question in ecology, conservation and island biogeography (Patiño et al., 2017). Extinct and introduced bird species can have distinct functional roles, and therefore some functions once performed by extinct birds may have disappeared from some islands (Sobral et al., 2016). However, it remains unclear how species extinctions and introductions translate into functional changes at the assemblage level, largely due to the difficulty in linking taxonomic and functional diversity.

Functional diversity is usually measured using species traits, under the assumption that these correlate to function (Cadotte et al., 2011). A simple but seldom used measurement of functional diversity change is to assess functional composition, obtained from the distribution of trait frequency based on the losses/gains of individual traits (Boyer & Jetz, 2014). However, in the past two decades, a multitude of mathematical approaches have been developed to estimate and visualize the functional diversity of assemblages (Cadotte et al., 2011; Mammola et al., 2020). These often follow the Hutchinsonian niche concept (Hutchinson 1957), relying on the position of species or individuals within a multidimensional space. Among these, the convex hull hypervolume is one of the most used, despite some important limitations, such as the assumption that the multidimensional space is homogenously occupied, making it extremely sensitive to outliers (Mammola & Cardoso, 2021; Mammola et al., 2020). To overcome this limitation, new methods have used probabilistic hypervolumes (Blonder et al., 2018), of which the most popular uses high-dimensional kernel density estimations to delineate the shape and volume of the multidimensional space (Carvalho & Cardoso 2020; Mammola & Cardoso, 2021).

Oceanic islands have long been used as living laboratories to understand ecological and evolutionary patterns and processes (Whittaker et al., 2017). Their discreteness, small size, simplified communities, unique biodiversity, and often recent human influence make them remarkably useful to study the impacts of human activities and explore promising conservation strategies (Russell & Kueffer, 2019). Our study aims to quantify the consequences of the species compositional changes promoted by extinctions and introductions on the functional diversity and composition of island bird assemblages. Focusing on 74 oceanic islands, we ask whether the functional diversity gained by introduced species has offset the reduction in diversity caused by the loss of native species. By adopting a functional perspective, we hope to gain valuable insights into the ecology of island bird assemblages and thus understand how to maintain their remaining functional diversity.

## 2. Methods

### 2.1. Island selection

We focused on the world’s biggest oceanic islands, those larger than 100 km^2^. From an initial list of 87 islands (Weigelt et al., 2015), we excluded 13 for which we were unable to obtain a species checklist or that do not have terrestrial or freshwater breeding bird species (see below and Table S1).

### 2.2. Bird species database

We compiled a list of known breeding bird species for each island, including extinct, extirpated and established introduced species, following the taxonomy used by Birdlife International (Handbook of the Birds of the World & BirdLife International, 2018). Given our focus on the temporal changes in species composition within islands, we also included island-level extirpations. For simplicity, we use the terms ‘extinction’ and ‘extinct’ for both global and local extinctions.

We excluded marine birds, non-breeding migrants, occasional breeders, vagrant and accidental species, and focus on regularly breeding terrestrial and freshwater species, since these are the most dependent on island resources and also have particularly high rates of extinction and introduction on oceanic islands (del Hoyo et al., 2014).

To obtain a complete list of species for each island, we first gathered checklists of extant birds from *Avibase* (Lepage, 2018). We then removed non-breeding species and flagged introduced species, based on *HBW Alive* (del Hoyo et al., 2014) and distribution maps in the *IUCN Red List of Threatened Species* (IUCN, 2020; Fig. S1). In addition, we revised information based on regional field guides (Table S2), which often include finer details on the status of each species in each island (e.g. breeding, introduced, extinct).

We used the *Global Avian Invasions Atlas* (Dyer et al., 2017a) as an additional source of information on introductions, and clarified inconsistencies by further exploring the literature (Table S3).

We only considered as extinct the native species classified as Extinct and Extinct in the Wild in the *IUCN Red List of Threatened Species* (IUCN, 2020), and as Extinct or Extirpated in *Avibase* (Lepage, 2018). To improve the list of extinct species, we analysed specific literature (Hume, 2017; Paleobiology Database, 2018; Fig. S2), and thoroughly reviewed extinction records for each target island (Table S4). Species classified as Probably Extinct in the literature (especially in Hume, 2017) and Critically Endangered – Probably Extinct in the *IUCN Red List of Threatened Species* were carefully analysed and considered extinct only when the *IUCN Red List of Threatened Species* supported this claim. We only included extinct taxa if these had been identified to species level, which is often not possible from fossil or historical records.

### 2.3. Bird species traits

For each species, we gathered information on body mass, foraging time, diet, foraging strata, volancy and habitat (Table S5). These traits are commonly used in studies evaluating bird functional diversity and summarising the effects of species on ecological processes and on responses of communities to environmental change (Boyer, 2008; Luck et al., 2012; Sobral et al., 2016).

For extant species, our main source of information regarding average body mass, foraging time (‘diurnal’ or ‘nocturnal’), diet and foraging strata was the *EltonTraits* database (Wilman et al., 2014). For the 40 (out of 617) species missing from this database, we inferred traits from the closest species in the genus (Table S6). We treated average body mass both as a continuous variable, and as an ordinal trait, based on the 20-quantiles categories: ‘very small’ (<16.422g); ‘small’ (16.422 - 37.240g); ‘medium’ (37.240 - 98.544g); ‘large’ (98.544 - 327.370g); and ‘very large’ (>327.370g). Regarding diet, we converted the information on the relative importance of each diet class in *EltonTraits* into six mutually exclusive binary classes: ‘granivore’ if > 50% seeds (class ‘Seed’ in *EltonTraits*); ‘herbivore’ if 50% plants (class ‘PlantO’); ‘frugivore’ if > 50% fruits and/or nectar (class ‘FruiNect’); ‘invertivore’ if 50% invertebrates (class ‘Invertebrate”); ‘carnivore’ if > 50% vertebrates, fish and/or carrion (class ‘VertFishScav’); and ‘omnivore’ if < 50% in any of the previous five classes. In addition, to capture the unique nectar-feeding strategy, we created one binary class, ‘nectivore’, identifying all species if class ‘Nect’ (nectar, pollen, plant exudates, and gums) in *EltonTraits* was higher than 30%. For foraging strata, we adapted the information on prevalence (i.e. time spent) from *EltonTraits* into seven binary classes: ‘ground’, ‘understory’, ‘midhigh’, ‘canopy’, and ‘aerial’ if prevalence > 50% in the respective strata; ‘water’ if the summed prevalence in ‘foraging below the water surface’ and ‘foraging on or just below the water surface’ > 50%; and ‘nonspecialized’ otherwise. Information about flight ability (volancy) was extracted directly from Sayol et al. (2020). Information about habitat was obtained from the first level of classification of the IUCN Habitats Classification Scheme (IUCN, 2020), combined into seven non-mutually exclusive binary classes: ‘forest’; ‘savannah’; ‘shrubland’; ‘grassland’; ‘wetlands’; ‘desert’; ‘artificial aquatic habitats’; ‘marine habitats’ (which combine IUCN categories ‘marine neritic’, ‘marine oceanic’, ‘marine intertidal’, and ‘marine coastal/supratidal’); ‘artificial terrestrial habitats’ (IUCN categories ‘artificial – terrestrial’ and ‘introduced vegetation’); and ‘rocky and subterranean habitats’ (IUCN categories ‘rocky areas’ and ‘caves & subterranean habitats’). The last three habitat classes combined IUCN habitat categories that had few and ecologically similar species, which we assumed to have similar responses to environmental variables.

For extinct bird species, we also used mostly *EltonTraits* to collect information on body mass, foraging time, diet and foraging strata (Wilman et al., 2014; Fig. S3). For missing species (96 out of 214) and traits, we explored additional references (Boyer, 2008; Sobral et al., 2016; Heinen et al., 2018; Crouch & Mason-Gamer, 2019; Case & Tarwater, 2020; IUCN, 2020; Sayol et al., 2020) (Fig. S3). Lastly, whenever information on a trait for a given species was still missing, we first attempted to derive it from descriptions of the species, or (if not possible) inferred it from the traits of the closest species in the genus (File S1).

### 2.4. Data analysis

Data processing and statistical analyses were done in R (v.4.0.4; R Core Team, 2021).

#### 2.4.1. Species compositional changes

We used species richness (alpha taxonomic diversity) to quantify the changes in species composition associated with bird species extinctions and introductions in each island. Then, we calculated: average loss as the average number of extinctions per island; average gain as the average number of introductions per island; and net change as the difference between gains and losses (including 95% confidence intervals based on all 74 studied islands). We also calculated changes in the overall number of extinct and introduced species (gamma diversity), and the net change across all islands.

#### 2.4.2. Effects of bird extinctions and introductions on functional composition

For each island and for each categorical trait (body mass, foraging time, diet, foraging strata, volancy and habitat), we assessed how extinctions and introductions affected functional composition, i.e. the prevalence of species associated with each trait class at the assemblage level. We did this by calculating, for each trait class in each island: ‘loss’, as the number of extinct species; ‘gain’, as the number of introduced species; and ‘net change’, as the difference between gain and loss. We then averaged results across islands, to obtain the average gain, loss and net change of species per island for each trait class, as well as the respective 95% confidence intervals. We calculated averages by considering only islands where the trait class was represented by at least one species, either extant or extinct.

The calculations above described used the absolute number of species gained or lost, but they were repeated using percentages of species gained or lost, to account for differences in the number of species between islands. Thus, for each island, we divided the number of species lost or gained associated with each trait class by the total number of species in the original avifauna (i.e. pre-extinctions, including extant native and extinct species, but not introduced species). This allowed us to verify if gain and loss were affected by island species richness.

For analysis of body mass as a continuous trait, we estimated, for each island, loss as the average body mass of extinct species, gain as the average body mass of introduced species, and net change as the difference between gain and loss. We then obtained average results and respective 95% confidence intervals by averaging losses, gains and net changes across islands.

#### 2.4.2. Effects of bird extinctions and introductions on functional diversity

For each island, we analysed how bird species extinctions and introductions affected functional diversity, using three measures based on probabilistic hypervolumes: functional richness (alpha functional diversity), functional originality of species and functional evenness (Fig. S4). To calculate these measures, we built a trait space from a matrix composed of all analysed species and 10 traits derived from those used in previous analyses (Table 1).

**Table 1.** Description of the 10 traits used to build the trait space.

| Trait | Type | Description |
| --- | --- | --- |
| Diurnal | Dichotomous | Diurnal (1), nocturnal (0) |
| Nectivore | Dichotomous | Nectivore (1), non-nectivore (0) |
| Water forager<sup>1</sup> | Dichotomous | Yes (1), no (0) |
| Forest specialist | Dichotomous | Forest specialist (1), non-forest specialist (0) |
| Wetland specialist | Dichotomous | Wetland specialist (1), non-wetland specialist (0) |
|  | Nominal | Granivore, herbivore, frugivore, invertivore, carnivore, omnivore |
| Average body mass | Quantitative | Natural log-transformed body mass |
| Habitat specialization | Quantitative | Number of suitable habitats listed by IUCN |
| Volancy | Ordinal | Flightless (1), weak flyer (2), volant (3) |
| Terrestrial foraging strata<sup>2</sup> | Ordinal | Ground (1), understory (2), midhigh (3), nonspecialized (3.5), canopy (4), aerial (5) |

**Note:** <sup>1</sup>Water forager and terrestrial foraging strata are not mutually exclusive, meaning that a species can be considered, for example, both water forager (1) and ground (1), as with most Anatidae species. <sup>2</sup>We considered nonspecialized birds, species that forage in most strata between ground and aerial, and thus attributed them the average value of 3.5.

We computed the pairwise functional distances between each pair of species using the Gower dissimilarity index, giving the same weight to each trait (range: 0 – 0.887), and then calculated the contribution of each trait to the resulting distance matrix, using respectively dist.ktab and kdist.cor in ade4 package (Table S7; Dray & Dufour, 2007). We analysed the distance matrix through a principal coordinate analysis (PCoA) with the Cailliez correction for negative eigenvalues to extract orthogonal axes for the hypervolume delineations, using the pcoa function in the ape package (Paradis et al., 2004). To construct the trait space, we retained the first eight PCoA axes, which cumulatively explained 81.3% of the total variation (Fig. S5).

In the probabilistic hypervolume, trait space was constructed using kernel density hypervolumes. These were approximated to a cloud of species-based stochastic points, which were positioned according to their traits in the multidimensional space. The kernel density hypervolumes were built using the Gaussian method with a 95% bandwidth so that hypervolumes represent 95% of the cloud density (Blonder et al., 2018). The functional richness of the assemblage is estimated as the volume of the hypervolume delineated by the cloud of stochastic points (Fig. S4; Mammola et al., 2020). This approach assumes a heterogeneous trait space, representing variations in point density within the multidimensional space. Thus, adding a species may decrease functional richness, namely if the species is close to the centre of the assemblage and decreases the average distance between points within the cloud.

For each island, we estimated the functional richness of the respective bird assemblage at two points in time: originally (i.e. including all native species, both extant and extinct); and presently (i.e. including extant native and introduced species). For this purpose, we used the kernel.alpha function in the BAT package (Cardoso et al., 2015; Mammola & Cardoso, 2021). Also for each island, we calculated net change in functional richness as the difference between present and original functional richness. Similarly, we evaluated the evenness of the total trait space for each island, considering original and present avifaunas, using kernel.evenness in BAT package (Fig. S4; Mammola & Cardoso, 2021), and calculated net change as the difference between the two. We then calculated average values of functional richness and evenness across islands, and respective 95% confidence intervals, for the original and the present avifaunas, and for the net change.

Finally, we evaluated the functional originality of each species, which is the average dissimilarity between the species and a sample of random points within the boundaries of the hypervolume. Within each island, the sum of values across all species is equal to one. We estimated originality based on a 0.01 fraction of random points, using the kernel.originality function in the BAT package (Fig. S4; Mammola & Cardoso, 2021). For each island, we calculated the average functional originality of the extinct species in the original avifauna, and of the introduced species in the present avifauna. From these values, we estimated average values of originality for extinct and for introduced species across all islands, and respective 95% confidence intervals.

## 3. Results

Our database included 759 species in 2709 island populations, distributed across 74 oceanic islands. Of these, 214 species and 280 populations were extinct, 172 species and 801 populations were introduced, and the remaining (445 species and 1628 populations) were extant natives (Table S1 and Fig. S6). Some species were introduced to an island but native to another, or extinct from one island while extant on another.

### 3.1. Species compositional changes revealed by species richness

Across all islands, there was a net decrease in the total number of species (gamma diversity), as there were more extinct than introduced species (Fig. 1a). However, average species richness per island (alpha diversity) experienced a positive net change, since the average number of introduced species on each island was higher than the number of extinct species (Fig. 1b and S8, Table S9).

**Figure 1.**
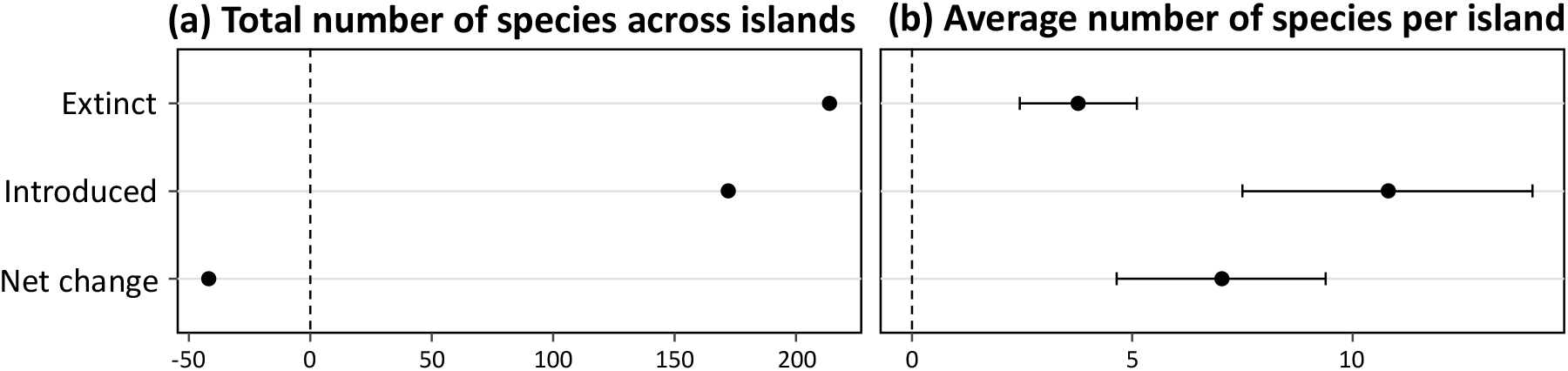
Total number of extinct and introduced species, and net change (introduced minus extinct) when considering (a) the total number of these species across all islands and (b) the average number of these species per island. In (b), the averages (circles) and 95% interval confidence estimates (horizontal bars) were obtained from values across each of the 74 islands. A negative net change indicates that the original avifauna tended to have a higher species richness than the present avifauna, whereas a positive net change indicates the opposite.

### 3.2. Effects of species compositional changes on functional composition

We found a positive average net change in the prevalence of 23 out of 35 traits (Fig. 2 and Table S8), meaning that, for each of those traits, the average number of introduced species per island associated with the trait was higher than that of extinct species. Conversely, net change was negative for seven traits and non-significant for five. Qualitatively similar results were obtained when correcting for islands’ species richness, with only three additional traits having a non-significant net change (nocturnal, invertivore and nectivore; Fig. S7). We thus focus on absolute numbers of introduced/extinct species.

**Figure 2.**
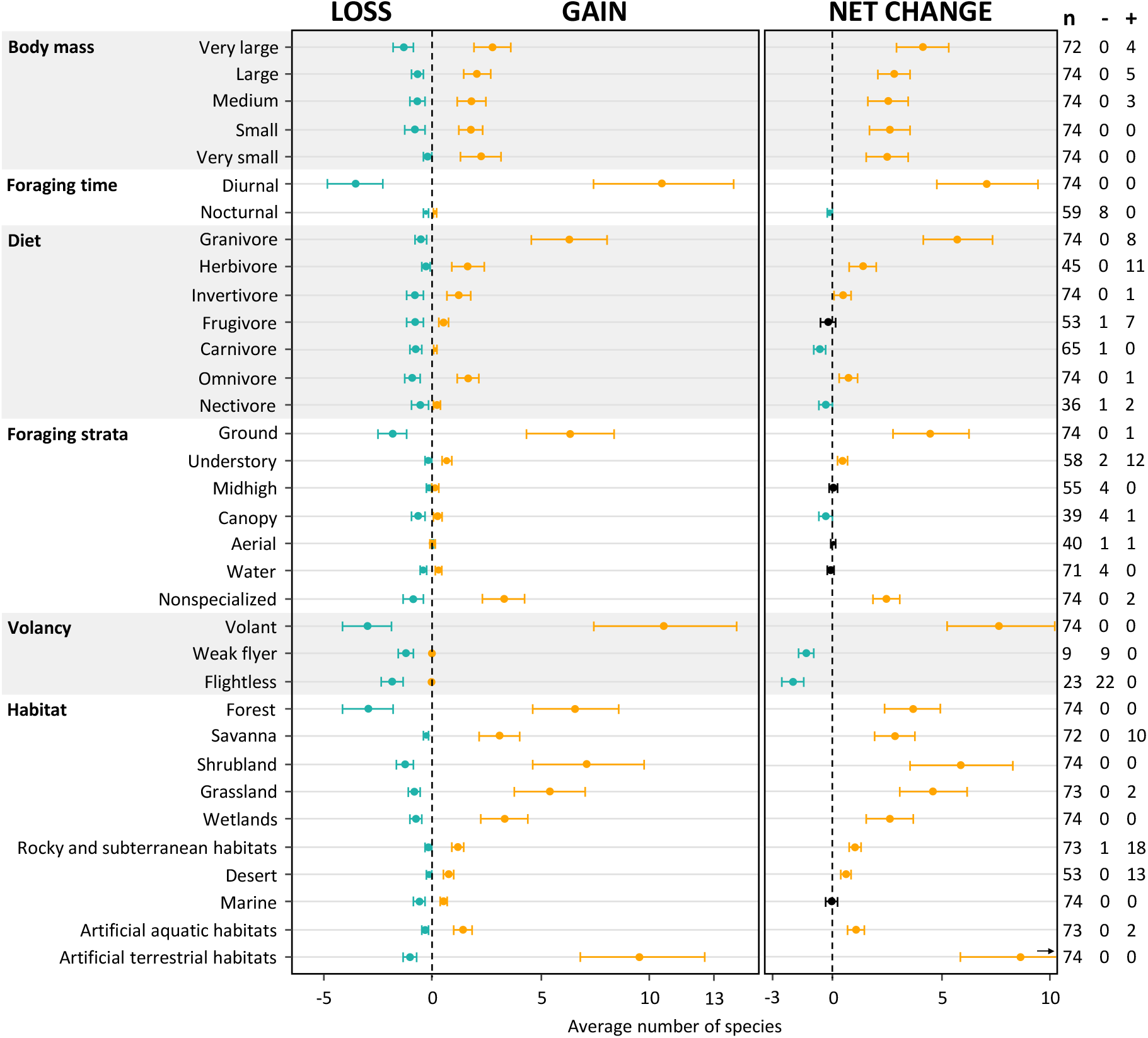
Effects of species compositional changes on island functional composition. For each trait class, we present the average number of species associated with the trait that were lost per island through extinctions (loss), gained through introductions (gain) and the difference between gain and loss (net change). Circles represent average values across islands, the horizontal bars the 95% confidence intervals. Non-significant values of net change (*p*-value > 0.05) are represented in black. Column ‘n’ represents the number of islands used in the calculations (i.e. with at least one species in the corresponding trait class), whereas columns ‘-’ and ‘+’ show respectively the number of islands that lost and gained species with a given trait.

We observed a positive net change across all classes of body mass (Fig. 2), meaning that more species were introduced than extinct in each size category. However, the average body mass of extinct species was higher than that of introduced species (natural log-transformed average body mass = 5.241g ± 0.284 > 4.513g ± 0.125, calculated across 52 and 73 islands, respectively), and there was a decrease in average body mass (−0.785 ± 0.348, calculated across 74 islands; Table S8).

We also found a positive net change in the prevalence of diurnal species, granivores, herbivores, invertivores, omnivores, volant species, ground, understory, nonspecialized foragers, and in species that occur in each habitat class, except marine habitats. In contrast, we found a negative net change in the prevalence of carnivores, nectivores, canopy foragers, weak flyers, and flightless species. The only introduced nocturnal bird species was the barn owl, *Tyto alba*, in all the Hawaiian Islands.

Within 4 out of 6 groups of traits, the class with the highest net change (very large body mass, diurnal foraging, ground foraging and volant species) had both the largest loss and the largest gain (Fig. 2). This suggests that these classes are, overall, the most susceptible to changes in species composition. In contrast, the class with the highest net change for both diet and habitat (granivore and occurring in artificial terrestrial habitats) had the greatest gain but not the greatest loss.

Introductions added on average more novel traits than those that had been lost by extinctions.

Indeed, for 19 trait classes, introduced species added novel traits to between 1 and 18 islands (column “+” in Fig. 2), while extinctions removed 12 traits on between 1 and 22 islands, (column “-” in Fig. 2). Weak flyers and flightless birds were particularly prone to extinction and have disappeared from almost all islands where they used to occur (weak flyers: 9/9 islands, flightless: 22/23 islands - only Campbell teal, *Anas nesiotis*, remains in Campbell Island).

### 3.3. Effects of species compositional changes on functional diversity

Despite the net positive change in average species richness per island, we found a net negative change in average functional richness (Fig. 3a and S8, Table S9). Introduced species with traits mostly similar to extant natives were responsible for reduced functional richness on 19 islands that had no extinctions.

**Figure 3.**
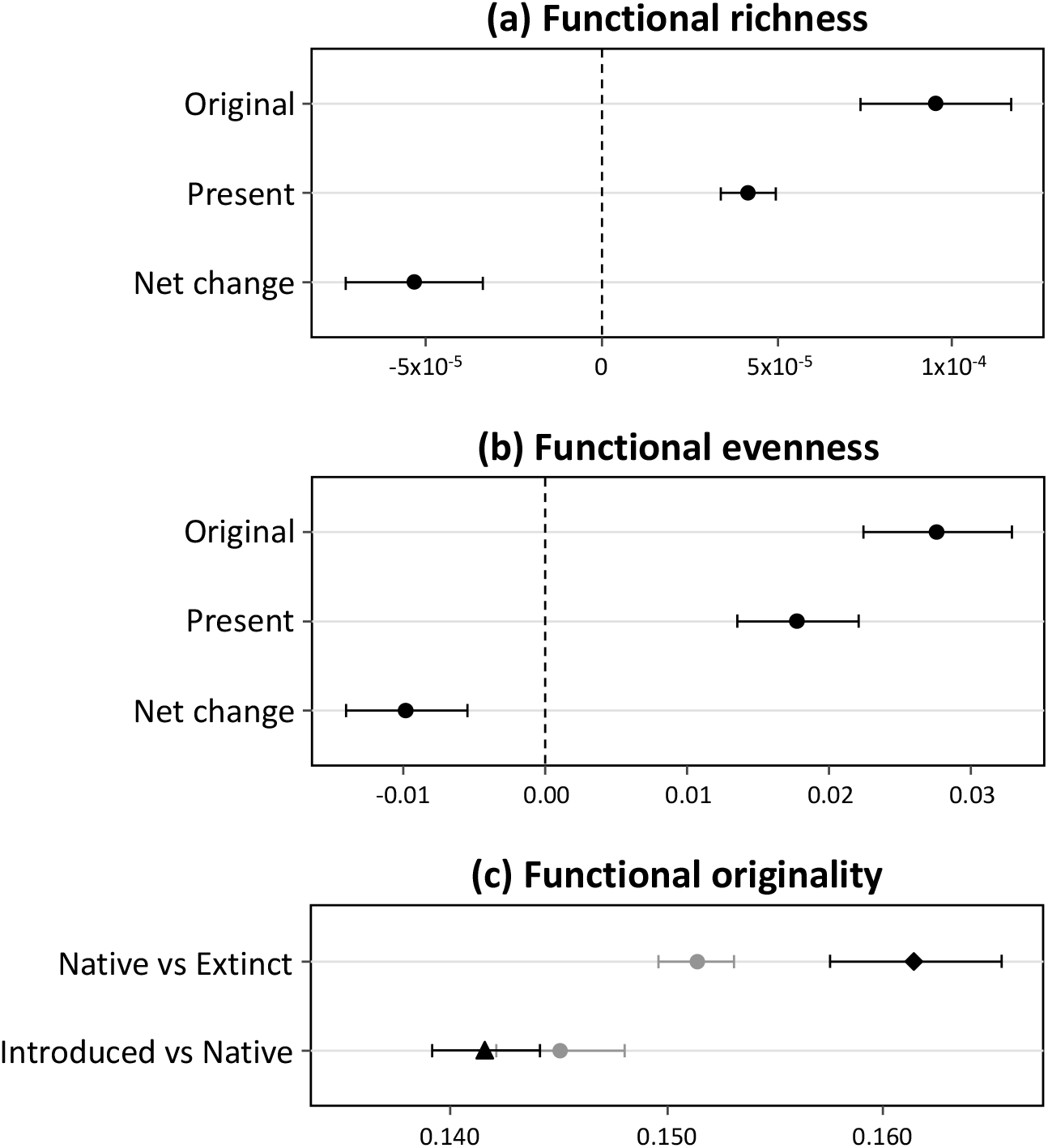
Effects of changes in the species composition of islands on three measures of functional diversity: (a) functional richness; (b) functional evenness; and (c) functional originality. Values presented are averages (circles) and 95% interval confidence estimates (horizontal bars) across islands. In (a) and (b), values correspond to the average volume of the trait space obtained from two probabilistic hypervolumes built for each of the 74 islands: one built with the species in original avifauna (extant native and extinct species), and another derived from the present avifauna (extant native and introduced species); net changes are the difference in volume between present and original: negative indicating a net loss in functional diversity; positive the opposite). In (c), we contrast the average functional originality of extant versus extinct native species (in the context of the original assemblages), and of introduced versus extant native species (in the context of the present assemblages). Average values were calculated, respectively, for the 52 islands with extinct species, the 73 with introduced species and the 74 with extant native species.

The species compositional changes also led to a negative net change in average functional evenness per island, indicating that the original avifauna was, on average, more evenly distributed across the trait space than the present avifauna (Fig. 3b and S8).

Compared with extant native species, the average functional originality of extinct species was significantly higher, whereas that of introduced species did not differ significantly (Fig. 3c and S9), meaning that extinct species have a more unique position within the trait space than either extant natives or introduced species.

## 4. Discussion

### 4.1. Increase in local species richness despite net losses across islands

We found an increase in the average number of bird species per oceanic island (alpha diversity), even though overall species richness decreased across all islands (gamma diversity; Fig. 1). This apparent paradox reflects the fact that a smaller overall number of species were introduced, but to multiple islands (Blackburn et al., 2009; Dyer et al., 2017b), than those that went extinct, often endemic to single islands (Boyer, 2008; Boyer & Jetz, 2014). This turnover in community composition associated with extinctions and introductions is likely increasing the similarity between island bird assemblages (i.e., lowering beta diversity, promoting biotic homogenization; McKinney & Lockwood, 1999). These findings are in line with previous studies (e.g. Sobral et al., 2016).

An incomplete knowledge of original island avifaunas creates uncertainty around these estimates of the magnitude of species compositional change. First, we are likely to underestimate the number of extinct species (Boehm & Cronk, 2021), given that new species keep being described (e.g. Rheindt et al., 2020). Second, it is not always clear which species are native or introduced (Essl et al., 2018). Furthermore, these results are a snapshot in time: the number of introduced species will very likely continue to increase in many islands (Seebens et al., 2017). Overall numbers of introductions may thus eventually surpass the overall number of extinctions.

### 4.2. Species compositional changes promoted changes in functional composition

We found evidence of significant changes in the functional composition of island bird assemblages, consistent with previous studies (e.g. Sax & Gaines, 2008). Overall, the net increase in average species richness per island was accompanied by positive net changes for 23 out of 35 traits, while negative net changes occurred only for seven (Fig. 2). Islands gained species of all size classes, as well as diurnal species, granivores, herbivores, invertivores, omnivores, volant species, ground foragers, understory foragers, nonspecialized forager species, and species that occur in all habitat classes but marine.

Very large bird species were simultaneously the biggest loss, gain and positive net change within body size classes, highlighting the volatility of these populations in oceanic islands. We also showed that the average body mass of extinct species was higher than that of introduced species, which led to a decrease in the average body mass of island bird species assemblages. This finding provides further support that large species are particularly prone to extinction (Boyer, 2008; Fromm & Meiri, 2021).

Regarding diet classes, the largest positive net gains in prevalence were by far of granivores, followed by herbivores, omnivores and invertivores. There was no significant net change for frugivores, while for carnivores and nectivores net changes were negative. Similar trends have already been described (Blackburn et al., 2009; Soares et al., 2021), and reflect a simplification of ecological networks: favouring lower positions in the trophic chain and unspecialized species, which are often better adapted to simplified anthropogenic landscapes, while hindering species that rely on more complex relationships, such as top positions in the trophic chain and nectarivory. These changes to island bird assemblages might disrupt well-established mutualistic plant-animal interactions and affect native plants, particularly through reduced pollination and seed dispersal (e.g. Caves et al., 2013; Carpenter et al., 2020). Herbivore birds introduced to islands that had no native browsers or grazers can greatly affect ecosystems, including by reducing food resources for pollinators and ultimately changing the phenotypic traits of plants related to pollination (e.g. flowering phenology, flower production, quantity and quality of nectar and pollen; Traveset & Richardson, 2006). Carnivore birds are more extinction-prone due to their high diet specificity (Şekercioğlu et al., 2004). Their loss can have serious negative consequences to ecosystems (Şekercioğlu, 2006), such as the increase of undesirable species and disease outbreaks if scavengers disappear, or the decline of guano and associated nutrients input if piscivores are loss (Şekercioğlu et al., 2004). Nectivore birds can also play a critical ecological role in the ecosystem, and their disappearance can have serious impacts on plant-bird mutualistic interactions, potentially impairing the future of insular native forests (Şekercioğlu et al., 2004; Boyer, 2008). This is particularly important in some island ecosystems that have few pollinators and many flowering plant species that depend exclusively on birds (Anderson et al., 2011).

Ground, understory, nonspecialized forager species had a positive net change in prevalence, while canopy foragers had a negative net change. This disappearance of birds adapted to forest foraging is a direct consequence of the extreme anthropogenic deforestation that occurred on many oceanic islands (Russell & Kueffer, 2019). Conversely, it also favoured the establishment of bird species that prefer open areas, which often have ground or unspecialized foraging strategies (Blackburn et al., 2009; Soares et al., 2021).

Flightless and weak flying birds can have important and sometimes irreplaceable ecological roles in key ecosystem functions (Boyer & Jetz, 2014), such as seed dispersal, pollination and herbivory (e.g. Carpenter et al., 2020) but they have been completely eradicated from almost all islands (Sayol et al., 2020; Fromm & Meiri, 2021). This proneness to extinction is linked to having evolved in the absence of mammalian predators, which were introduced to most islands (Milberg & Tyrberg, 1993; Russell & Kueffer, 2019). Introduced mammals sometimes occupy ecological niches similar to those of flightless birds, making it very difficult to re-establish populations of flight-impaired birds.

For all habitat classes, except marine, there was a positive net change in prevalence. The largest net gain was for species that occur in artificial terrestrial habitats, followed by other open habitats (shrubland and grassland). Even though there was a clear net gain in the prevalence of forest species, they were also the ones most subject to extinctions. The loss of forest-dependent birds due to the removal of native forests on oceanic islands is well known (e.g. Pimm et al., 2006; Hume, 2017). In the Hawaiian Islands, for example, hunting and destruction of lowland forest by Polynesians extinguished many endemic forest birds, long before European arrival (Olson & James, 1982).

Overall, although islands have gained more bird species than they have lost, the functional composition of their avifaunas has changed markedly, potentially with consequences to ecosystem functioning (e.g. Heinen et al., 2018).

### 4.3. More species but with common traits, resulting in decreased functional diversity

Even though the combined effect of bird extinctions and introductions was a net gain in average island species richness (Fig. 1b) and an increased prevalence of most traits (Fig. 2), counterintuitively this translated into a decrease in average island functional richness (Fig. 3a). This indicates that the introduced species tend to be functionally more similar to remaining native species than extinct species were, resulting in a more compact cloud of points in the multidimensional trait space. Accordingly, we also observed a decrease of assemblage functional evenness (Fig. 3b), and found that whereas extinct species were functionally more original than the remaining native ones, the reverse is true for introduced species (Fig. 3c). The non-random extinction and introduction of bird species was already known to impair the functional diversity of island bird assemblages (Boyer & Jetz, 2014) since introduced species do not compensate for the functional roles of extinct species (Sobral et al., 2016). Surprisingly, the overall trends of taxonomic and functional richness only coincided (both decreasing or increasing) in 10 out of 74 islands (13.5% - Fig. S7). In 57 islands (77%), functional richness decreased despite increased species richness, while in three islands (Socorro, Floreana and San Cristóbal) there was an increase of functional richness despite a decrease in species richness (4.1%).

Islands are well-known for their high levels of endemism, unique functional traits and peculiar evolutionary patterns (Whittaker et al., 2017; Russel & Kueffer, 2019). Unfortunately, this uniqueness also makes insular species prone to anthropogenic extinctions (Hume, 2017), and their functions more difficult to replace (Boyer & Jetz, 2014). Introduced species tend to have specific ecological niches, distinct from those of natives, and prefer human-modified landscapes (Lee et al., 2010; Soares et al., 2021), which does not favour functional diversity. However, bird trade is turning to Neotropical species (Dyer et al., 2017b) and that might affect the traits of future introduced bird species, being better functional replacements of extinct species or, most likely, having a great potential to outcompete native species and further push these assemblages towards a functional collapse (Soares et al., 2021).

### 4.4. Maintaining functional diversity

We showed that a gain of species does not necessarily imply a gain in functional diversity, illustrating why these two facets of biodiversity should be assessed simultaneously to understand the impacts of human activities on biodiversity and ecosystem functioning. This mismatch between taxonomic, functional and even phylogenetic diversity has been observed across multiple different taxa (Brum et al., 2017; Albouy et al., 2017), and has challenged the use of surrogates (Devictor et al., 2010). Traditionally, global conservation efforts focused on protecting species or sites that have high conservation value, inadvertently underrepresenting other facets of biodiversity, such as functional diversity (Cadotte & Tucker, 2018). The non-linear and often negative relationship, as in oceanic island birds, between taxonomic and functional diversity calls for the prioritized protection of functionally unique species to maintain functional diversity. Such a line of action is also key to ensure that functional redundancy is kept, since it allows maintaining ecosystem functions under species loss.

In order to maintain functional diversity, we must prevent further loss of native ecosystems because their functioning depends on complex and irreplaceable ecological interactions (Aslan et al., 2013; Carpenter et al., 2020; Carmona et al., 2021). We also must avoid new introductions, especially of species that might affect ecosystem functions, either through predation (Milberg & Tyrberg, 1993; Sax & Gaines, 2008; Loehle & Eschenbach, 2011), competition (Soares et al. 2021), or the disruption of mutualistic interactions (Caves et al., 2013; Carpenter et al., 2020). Lastly, we need to protect native species, focusing on those that have unique functional traits.

Functionally unique species have recently been considered key for effective conservation because they represent distinct ecological strategies and often have disproportional higher extinction risk (Griffin et al., 2020; Carmona et al., 2021). However, in order to preserve the global diversity of ecological strategies, conservation efforts have to integrate complementary metrics, such as functional richness and functional uniqueness at multiple scales (Cooke et al., 2020). Many shortcomings still impair this integration, notably the lack of trait and distribution data for most taxa. Nevertheless, our work provides a framework for capturing changes in functional diversity, using functional richness, functional evenness and species functional originality at multiple scales, which can be extended to other taxa and other drivers of biodiversity change.

## 6. Data accessibility statement

All data supporting the results in the paper are archived in figshare and should be available once the paper is accepted and published.

## 7. Acknowledgements

This work was funded by the Portuguese Government “Fundação para a Ciência e para a Tecnologia” (FCT/MCTES), through FCS’ PhD grant (PD/BD/140832/2018) and cE3c’s Unit funding (UIDB/00329/2020). We are grateful to Dr Ruben Heleno for his help to validate the Galápagos bird list.

## 9. Biosketch

**Filipa Coutinho Soares** has an MSc in Conservation Biology by the Faculty of Sciences of the University of Lisbon. She has studied the synergistic effects of land-use and exotic species on the endemic-rich bird assemblage of São Tomé Island (Gulf of Guinea, Central Africa). Currently, she is a PhD candidate at the same faculty, studying the consequences of bird species extinctions and introductions on native biodiversity across oceanic islands worldwide.

